# Transcriptional analysis in bacteriophage Fc02 of *Pseudomonas aeruginosa* revealed two overlapping genes with exclusion activity

**DOI:** 10.1101/2022.11.17.516636

**Authors:** Irais Ramírez-Sánchez, Marco Magos-Castro, Gabriel Guarneros

## Abstract

Little is known about the gene expression program during transition from lysogenic to lytic cycles of temperate bacteriophages in *Pseudomonas aeruginosa*. To investigate this issue, we developed a thermo-sensitive repressor mutant in a lysogen and analyzed the phage transcriptional program by strand-specific RNA-Seq before and after thermo-induction. As expected, the repressor gene located on the phage DNA forward strand, is transcribed in the lysogen at the permissive temperature of 30°C. Upstream the promoter gene, we noticed the presence of two overlapped ORFs apparently in the same transcript, one ORF is a gene that encodes a protein of 7.9 kDa mediating exclusion of various super-infecting phages. The other ORF, placed in an alternate reading frame, with a possible AUG initiation codon at 25 n downstream the AUG of the first gene, is expected to encode a 20.7 kDa polypeptide of yet unknown function. Upon lifting repression at 40°C, starts transcription of an operon, involved in the lytic cycle from a promoter on the reverse phage DNA strand. The first gene in the operon is a homolog of the antirepresor *ner*, a common gene in the lysis-lysogeny regulation region of other phages. Interestingly, the next gene after *ner* is gene10 that on the reverse strand, overlaps the overlapped gene *olg1* on the forward strand. Curiously, gene 10 expression also shows superinfection exclusion. Strand-specific RNA-Seq also has uncover the transcription succession of gene modules expressed during the phage lytic stage.

## 1 Introduction

During lysogeny the resident bacteriophage, named prophage, renders the lysogenic cells immune to secondary infection by bacteriophages of the same type as the prophage. This immunity is operated by the repressor, a protein directed by the prophage that turns-off its lytic functions as well as those of the incoming homologous phage (Salmon et al., 2000). The prophages also may prevent infection of their hosts by a wider range of other phages different from the resident phage. This event named superinfection exclusion, or simply exclusion, is usually acted by proteins encoded in the prophage genome (Ali et al., 2014, Hofer et al., 1995, McGrath et al., 2002, Cumby et al., 2012, Owen et al., 2021). Strain Ps33 of *P. aeruginosa* lysogenic for phage Ps56 is refractory to infection by a collection hetero-immune test phages that readily infect the non-lysogenic parental strain. It has been suggested that the concerted action of two genes, named 9 and 10, of phage Ps56 was responsible for the exclusion(Carballo-Ontiveros et al., 2020). It also has been documented that filamentous phages Pf inserted in the bacterial chromosome of *P. aeruginosa* confer exclusion by expressing a protein that interacts with PilC (Schmidt et al., 2022).

It may also occur interference against superinfection by secondary infecting phages in bacteria already engaged to lysis by a previous infection with virulent phages(Shi et al., 2020). In *P. aeruginosa* it has been documented cases of genes encoding superinfection exclusion by both temperate and virulent phages(Bondy-Denomy et al., 2016, Xuan et al., 2022, Schmidt et al., 2022, Tsao et al., 2018).

The phage Fc02 belongs to a family whose archetype is phage B3. The DNA of phage B3 has been sequenced and annotated (Braid et al., 2004). Two possible overlapped promoters were identified in the region that controls the phage lysis-lysogeny responses. Promoter pC2 in the direction of the immunity repressor gene *repc* in the DNA forward strand, and pE possibly transcribing the antirepresor gene *ner* in the reverse strand. This promoter configuration is prevalent in other phages(Jakhetia and Verma, 2015, Ranquet et al., 2005, Salmon et al., 2000, Wu et al., 2021). A third possible promoter was also located between ORFs 15 and 16 in the reverse strand of B3 DNA transcribing away from *ner* (Braid et al., 2004). Also, a third possible promoter transcribing in the repressor gene direction was described in Mu genome (Goosen et al., 1984).

The application of massive techniques of nucleic acids and protein sequencing as well as of bioinformatics algorithms in biology has revealed the presence of overlapped genes in genomes more frequently than previously assessed(Wright et al., 2022). The configuration of overlapped open reading frames (ORFs) and their expression signals is variable; they could be overlapped partially or totally, in one direction on the same DNA strand, or in opposite strands and directions(Rogozin et al., 2002, Schlub and Holmes, 2020).

In bacteria and their bacteriophages numerous cases of extensive overlapped genes have been described(Zehentner et al., 2020, Muñoz-Baena and Poon, 2022, Scherbakov and Garber, 2000). However, frequently the evidence provided is partial, the function of one or both overlapped genes is unknown or is not clear whether the corresponding mRNA is transcribed and translated into protein. More frequently the regulation regime of the proteins expressed, and their functions are unknown (Kreitmeier et al., 2022). The historic case of overlapped genes in phage φX174 has been updated recently (Sanger et al., 1977). Data from modified genomes led to conclude that gene overlapping in the phage genome is essential for the correct expression of proteins and DNA replication (Wright et al., 2020).

In the present work we have identified a cluster of three overlapped ORFs in the regulatory region that controls the lysogeny and lysis stages of phage Fc02. Two of the ORFs, *olg1* and olg2 are overlapped in the same direction on the forward DNA strand but in different reading frames. The third ORF, named gene 10, is overlapped extensively to the *olg1* and *olg2* pair on the DNA reverse strand. Here we present evidence that *olg1-olg2* are possibly transcribed from a promoter sequence identified right upstream from *olg1-olg2*, active during lysogeny. On the opposite strand gene 10 is part of an operon probably transcribed from a promoter located upstream the antirepresor gene *ner*, early in the lytic stage in the phage development. We show that *olg1* encodes a 7.9 kDa polypeptide that excludes other infecting phages through lysogeny. The olg2 ORF is expected to encode a 20.7 kDa polypeptide without apparent exclusion function.

## 2 Material and Methods

### 2.1 Bacterial strains and phages

Phage Fc02 was isolated from a collection of clinical strains of *P.aeruginosa* (Castañeda-Montes et al., 2018). The strains used in this experiment were *Escherichia coli* DH5α and *P.aeruginosa* PAO1(Sepúlveda-Robles et al., 2012). Strains were cultured in LB (Luria-Bertani) broth with shaking at 200 rpm at 30°C and 40°C. The information of the plasmids used in this work is given in Table 1. When required the following concentrations of antibiotics were used: ampicillin (*E. coli* 100μg/ml) and gentamicin (*E. coli* 15μg/ml, *P. aeruginosa* 50μg/ml)

**Table 1.**
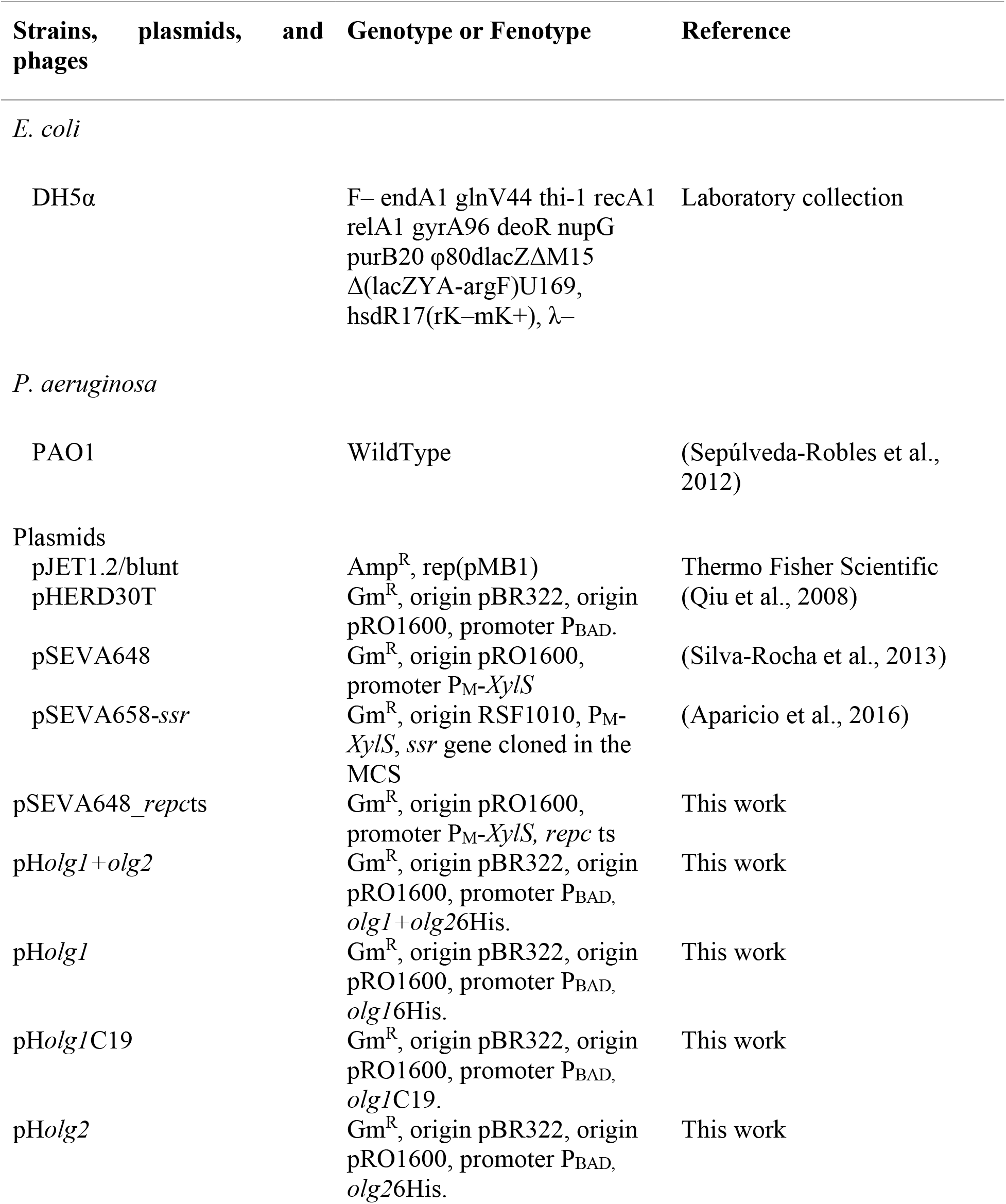
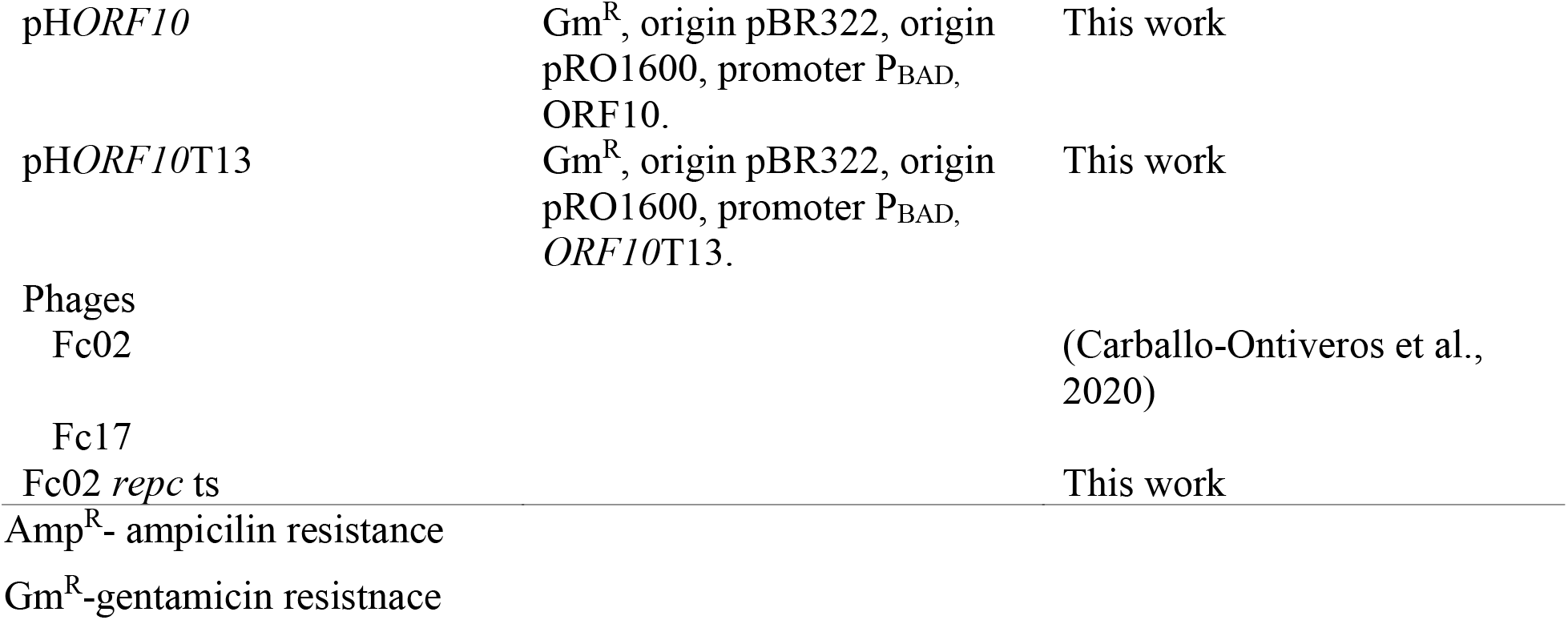
Strains, plasmids, and phages.

### 2.2 Random mutagenesis

An exponentially growing culture of PAO1(Fc02) was mutagenized with Nitrosoguanidine (NSG) as described in(Vogel et al., 1991) with modifications. Briefly, 7 ml of bacterial culture at an OD_600_ of 0.3 were incubated 30 min at 30°C with 50 μg/ml of NSG. Cultures were diluted 100-fold and recovered overnight in aeration at 30°C. The bacterial culture was centrifuged at 8,000 rpm for 5 min at 40°C to pellet the cells. The resultant pellet was washed 5 times with 5 ml of LB broth. Finally, the pellet was resuspended with 10ml of fresh LB broth and incubated with aeration for 3 hours at 40°C. The lysogen culture was centrifuged at 10,000 rpm for 10 min. The supernatant was treated with Chloroform to remove bacterial remnants and a virion suspension was carefully recovered (Echeverría-Vega et al., 2019) and plated on PAO1 lawns at 40°C. Clear plaques were isolated and plated on PAO1 lawns at 30°C. The plaques clear at 40°C and turbid at 30°C were isolated, and the lysogen was recovered and isolated from the turbid plaque center at 30°C.

### 2.3 Construction of the *repc* ts lysogen

The PAO1(Fc02 *repc* ts) lysogen was obtained with a modified version of the protocol described in (Rice et al., 2009). The point mutation G191A was introduced into the lysogen by homologous recombination. The repressor gene was cloned from the Fc02 ts mutant obtained with NSG, into the vector pJET1.2/blunt, used as suicide vector. Following the cloning method (ThermoFisher CloneJET PCR Cloning Kit), the pJET suicide vector was purified using the Jena Bioscience Plasmid Mini-Prep Kit and electroporated into PAO1(Fc02) and into PAO1(Fc02) harboring pSEVA658_PPβ (Aparicio et al., 2016). One milliliter of LB medium was added immediately after electroporation and cells were transferred to a 1.5 ml tube and incubated for 4 h at 30°C with shaking at 200 rpm. The culture was centrifuged at 10,000 rpm for 5 min, the supernatant was discarded, and the pellet washed twice with fresh LB broth. The pellet was resuspended in 1 ml of fresh LB and incubated with shaking at 40°C for 3 h. Culture was centrifuged at 10,000 rpm for 10 min and supernatant was collected and treated with chloroform. Finally, the phage in the supernatant was plated on PAO1 loans at 40°C to evaluate the thermo-sensitive phenotype. Independent mutants were purified and saved.

### 2.4 Lysogen growth-curve

A 50 ml culture of PAO1(Fc02 *repc* ts) was grown on LB broth with 200 rpm shaking at 30°C until OD_600_ 0.3, 2 ml of preheated medium was added to rapidly reach 40°C up-shifted temperature. Samples of 1ml were collected 5,10, 20 and 40 min after increasing the temperature, the phages at each time point including point 0, were diluted and plaques were evaluated in a PAO1 lawn. PFU/ml was calculated and plotted against time.

### 2.5 RNA extraction and sequencing

RNA was extracted from a modified protocol from(Blasdel et al., 2018). Fifty milliliters of LB broth were inoculated with 0.5 ml of overnight culture of PAO1 and PAO1(Fc02 *repc* ts) and incubated with shaking at 200 rpm at 30°C until an optical density at 600nm (OD_600_) of 0.2-0.3 was reached; the cultures were collected at 0, 5, 10, 20 and 40 minutes after thermal shifting to 40°C. Cell suspensions were treated with cold Stop Solution (95% EtOH, 5% phenol) and placed on ice until samples were pelleted by centrifugation for 20 min at 40°C at 8,000 rpm. After removing the supernatant, cells were resuspended in lysis buffer and total RNA was isolated with hot phenol extraction protocol, followed by ethanol precipitation. DnaseI (Thermo Fisher Scientific) treatment was done, and hot phenol extraction was repeated to remove any residual DnaseI, the integrity of the RNA was evaluated on a 1% agarose gel and RNA concentration was determined by NanDrop2000 (Thermo Fisher Scientific). To verify the absence of residual gDNA we performed a PCR on the RNA samples. RNA, samples were sent to GENEWIZ (New Jersey, USA) for Strand-Specific RNA sequencing using Illumina HiSeq platform and 150-bp paired-end reads.

### 2.6 Bioinformatic analysis

For the analysis of the 3D-structures prediction we used the software I-TASSER(Yang et al., 2015), the resulting model was compared to lamda CI repressor crystal structure under accession number PDB3J50; alignment of the two structures was done in ChimeraX. For the search of transcriptional regulators, we used BProm, SAPPHIRE, PhagePromoter, Promoter Prediction by Neural Network, FindTerm and ARNold(Solovyev and Salamov, 2013, Coppens and Lavigne, 2020, Reese, 2001, Naville et al., 2011). The fasta sequence was uploaded in ORF Finder for ORF search (Stothard, 2000).The predicted ORFs were analyzed using BLASTp against non-redundant protein databases (Altschul et al., 1997). Illumina fastq files were used as input in TRIMMOMATIC(Bolger et al., 2014) and FastQC. For the alignment we used HISAT2(Kim et al., 2019) with references genomes Fc02 (MH719189.1) and PAO1 (NC_002516.2). Coverage plots were created using custom scripts in R. Table counts were created from the counts of the Fc02 and PAO1 genes, and we used Deseq2 software to calculate differentially expressed genes between the different time groups (Love et al., 2014). Genes with a normalized counts fold change (FC) ≥1.5 and ≤-1.5 and a *p*-value ≤1×10^-5^ were considered differentially transcribed. The Venn diagrams were obtained using the jvenn viewer (Bardou et al., 2014).

### 2.7 Translatability of genes and exclusion activity evaluation

The genes *repc*, *olg*1 and 10 were cloned with EcoRI and HindIII into plasmid pHER30T in a 2-step modification(Qiu et al., 2008) (see primers in Table 2), and electroporated into PAO1. The exclusion phenotype of the ORF’s overexpression was evaluated by infection assay with test temperate phages on agar plates supplemented with gentamicin (Carballo-Ontiveros et al., 2020). Protein expression was performed growing the transformed bacteria in LB medium supplemented with gentamicin to OD_600_ 0.3 followed by induction with 0.3% of arabinose for 3 h. The genes *olg1* and *olg2* expression was carried out adding a 6 Histidine Tag at the 3’ ends, protein enrichment and purification was performed as in(Bertani et al., 1999) using a Ni^2+^ nitriloacetic acid metal-affinity column according to the manufacturer’s instructions (QIAGEN). Proteins were resolved by tricine-SDS-PAGE (Schägger, 2006) stained with Coomassie brilliant blue R-250 after electrophoresis or alternatively, proteins were transferred onto a nitrocellulose membrane. The membrane was subjected to Western blot analysis using a polyclonal antibody against the 6Histidine tag (Sigma-Aldrich). After incubation with the second anti-mouse antibody (Invitrogen) the purified protein was detected with SuperSignal^™^ West Femto (Thermo Fisher Scientific) and scanned in the LI-COR C-DiGit.

**Table 2.**
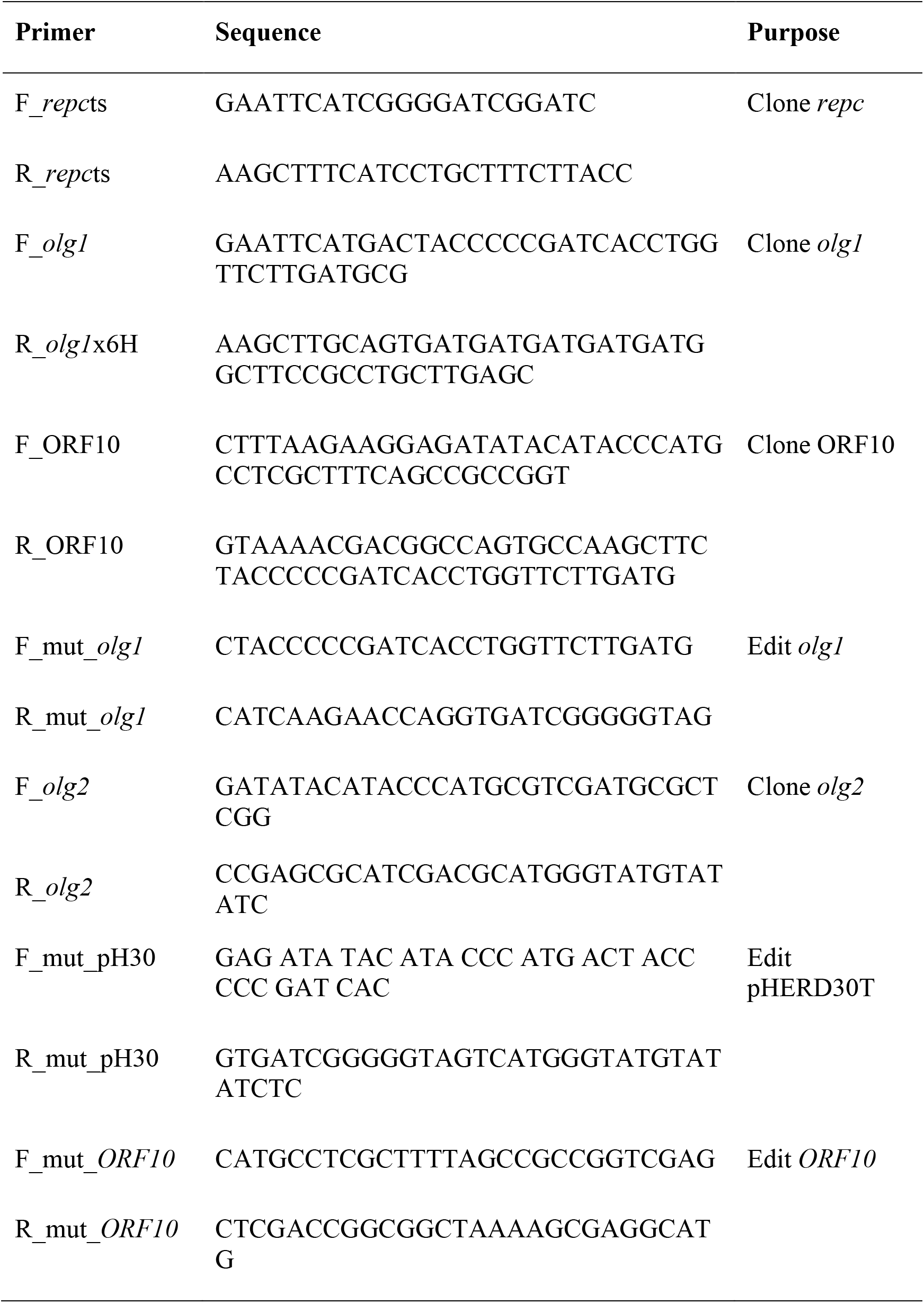
List of primers used to clone Fc02 ORF’s in pJET1.2/blunt, pHERD30T and pSEVA648.

## 3 Results

### 3.1 Generation and characterization of bacteriophage Fc02 thermo-sensitive repressor mutant

We approached the study of the transcriptional transition from lysogeny to lytic state of Fc02 by creating a thermo-inducible prophage variant(Liu et al., 2013). A culture of the lysogenic strain PAO1(Fc02) was mutagenized with nitrosoguanidine, up shifted from 30°C to 40°C and the supernatant plated on a PAO1 lawn on agar plates to select for clear plaques at 40°C that yielded turbid plaques at 30°C, the expected phenotype for the searched mutant. In one candidate we confirmed the presence of a mutation in the repressor gene by sequence and function. The position of the putative repressor gene in the genome of Fc02 was inferred by informatics assisted homology with the annotated sequence with lambda repressor (Carballo-Ontiveros et al., 2020). Sequencing of the presumed repressor gene from the isolated candidate showed the unique mutation G64E (Figure 1A and Supplementary Figure1). We cloned the mutant repressor gene in a suicidal vector to recombine the mutation into a wild type lysogen PAO1(Fc02) (see Material and Methods). Alignment of an informatic 3-dimensional model of Fc02 repressor protein, RepC, compared to structure solved for CI lambda repressor showed that the structures overlap closely in the HTH domain (Stayrook et al., 2008). Accordingly, the amino acid substitution in the *repc* ts mutant occurred in the second amino acid beyond the a5 helix of the HTH motif. The thermal sensitivity of the mutated repressor was functionally confirmed by phenotype of PAO1 bacteria transformed with a construct harboring the mutant repressor gene. This transformant was refractory to infection at 30°C but produced plaques upon infection with a Fc02 phage stock at 40°C (Figure 1B).

**Figure 1.**
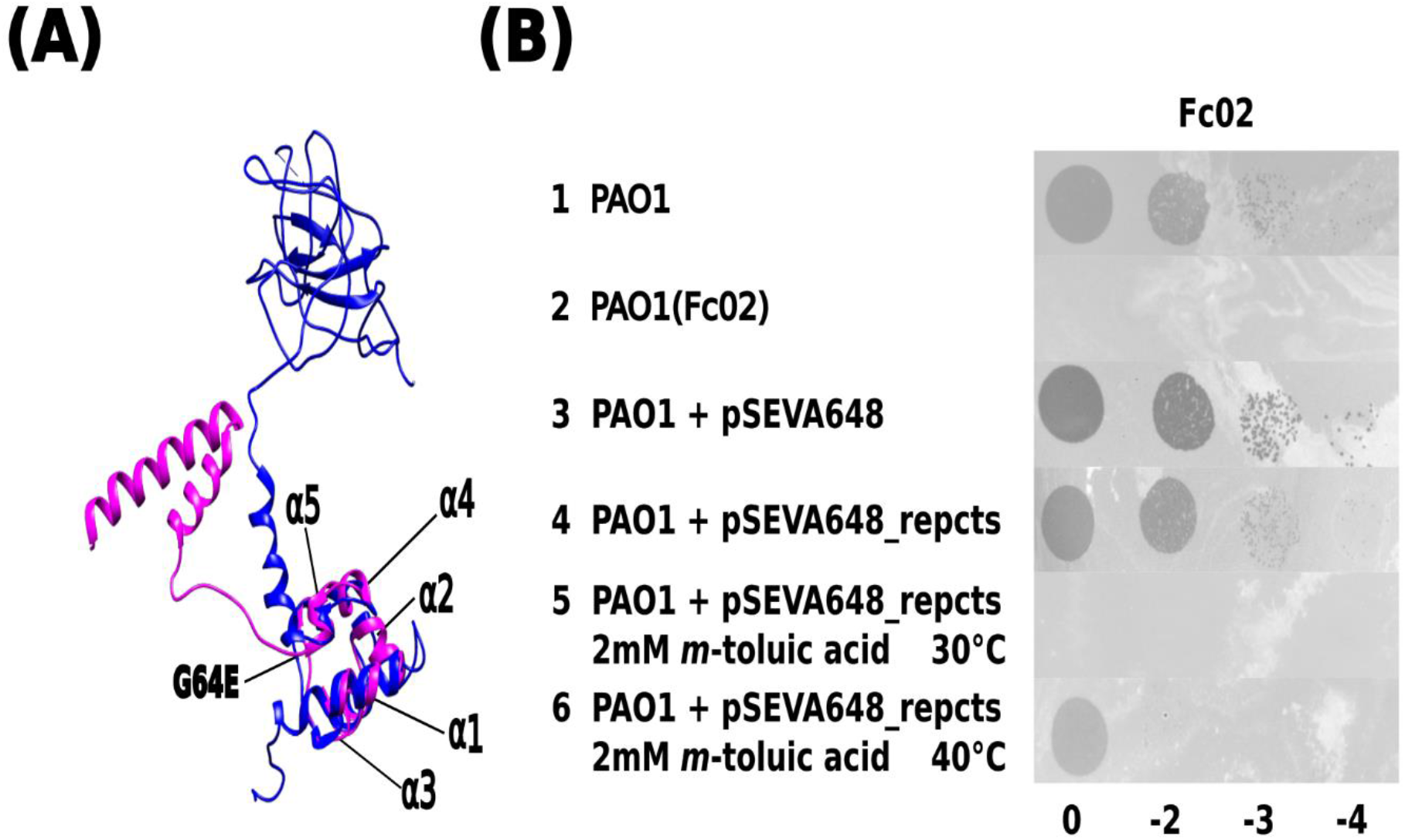
Identification of phage Fc02 immunity repressor gene *repc* and thermolability of *repc* ts mutant. A. Model of the structural homology of phage lambda CI repressor X-ray crystal resolved structure(Stayrook et al., 2008) (blue) and the predicted 3D structure of Fc02 ORF12 putative protein RepC (magenta). Notice the N-terminal domains overlapping of the proteins. The *repc* ts mutation G64E was located just beyond the last carboxy-side α-helix of the HTH motif of the putative protein. B. Drop dilutions lysis pattern of phage Fc02 on the indicated strains and temperatures. The phage stock decimal dilutions used in each column are indicated at the bottom of the panel. Notice that Fc02 infects the uninduced transformant (row 4), as it does wild type strain (row 1) but is unable to infect the *m*-toluic induced transformant at 30°C (row 5) and the PAO1(Fc02) lysogen (row 2).

### 3.2 Phage Fc02 transcription profile upon thermo-induction of a *P. aeruginosa* lysogen

To investigate the pattern of phage gene transcription during phage development, we designed an experiment of phage thermo-induction of the lysogen PAO1(Fc02 *repc* ts) (Figure 2A) and evaluate phage RNA at the appropriate time intervals. It has been documented that long before the phage progeny appears in the medium, phage genome transcription in the cell has ended (Wang, 2006). This seems to be the case for induction of Fc02 (Figure 2B). The last point analyzed for the phage DNA transcription was at 40 min of thermo-induction, a few minutes ahead the time of phage release to the medium and the decrease in the number of viable cells (Supplementary Figure 2).

**Figure 2.**
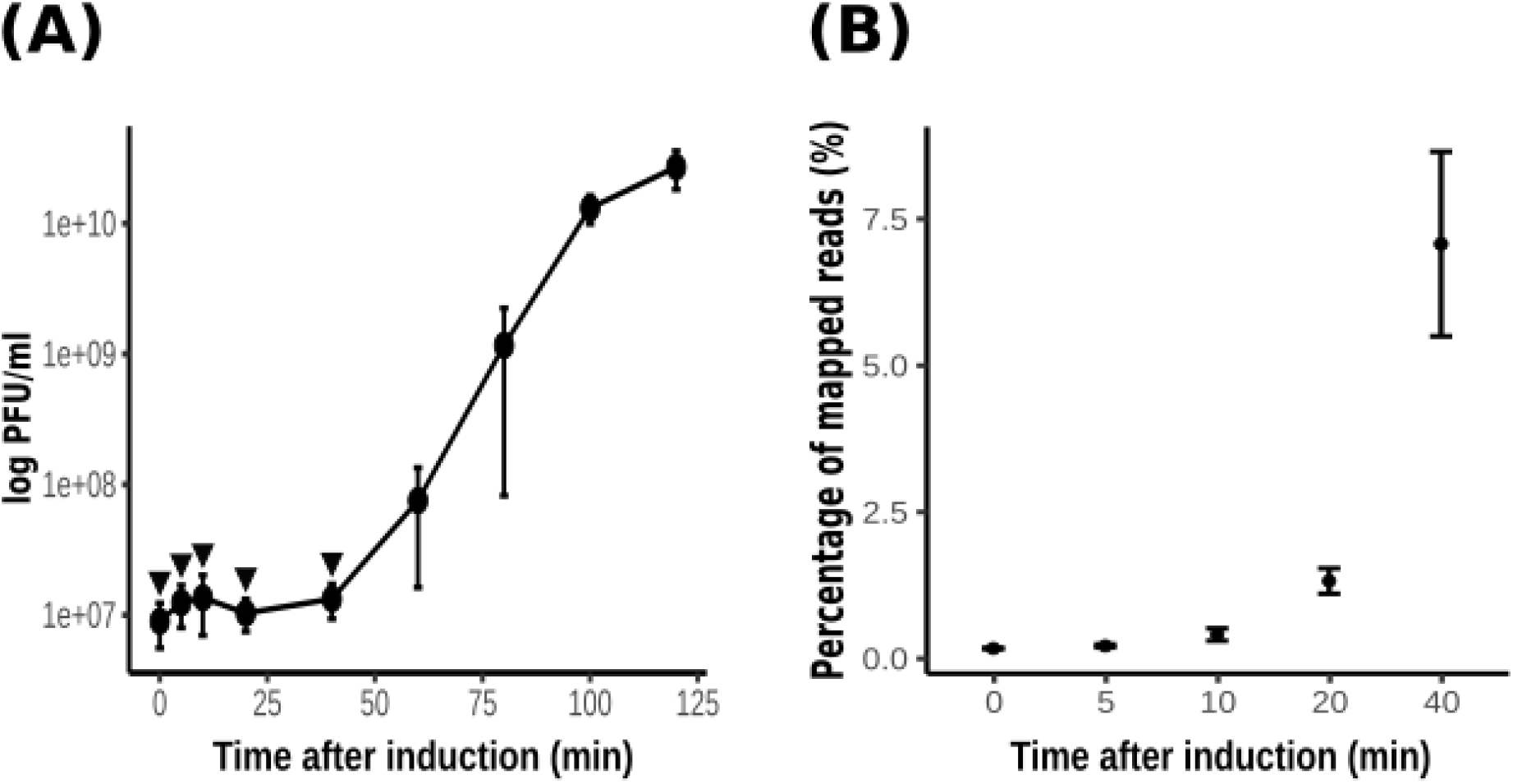
Thermo-induction of lysogen PAO1(Fc02 *repc* ts). A. Time course of the phage growth as measured by plaque forming units (pfu) on a lawn of strain PAO1. The lysogen was grown at 30°C to log phase, at t=0 the culture was up shifted and incubated at 40°C for 2h. Samples were drawn at the indicated times (filled circles) to quantify pfus. B. Samples (arrowheads in A) were drawn at the indicated times to be processed for strand-specific RNA-Seq. Most of the reads mapped to the bacterial sequences, but the percent of reads that mapped to the phage genome are plotted against time of thermo-induction. The average of the reads for the three biological replicas at each time was plotted and error bars indicate standard deviation

The wild type lysogen PAO1(Fc02) growing on liquid medium at 30°C did not significantly increase the release of infective phage upon shift to 40°C (Supplementary Figure 3). Therefore, no RNA-Seq sample was gathered for this strain. To investigate the gene transcription program of Fc02 in a lysogen in transition to the lytic mode, a lysogen growing at 30°C was up shifted to 40°C and samples were taken at various times (Figure 3). The profile shows that only forward strand mapped sequences in the regulation region. The transcripts include the *repc* gene as expected for a lysogen. The transcribed region of about 1900 nucleotides also includes ORFs in the DNA segments before and after de *repc* gene. The ORF downstream the repressor gene is the accessory gene, *e*4 without assigned function. This gene is not present in all the genomes of beetreviruses (Carballo-Ontiveros et al., 2020). Interestingly upstream the repressor gene two overlapped ORFs were identified on the forward strand of the phage DNA. The ORFs, named *olg1* and *olg2*, appear be transcribed from a promoter sequence, named pC1, located right upstream *olg1*. Presumably, the *olg1* and *olg2* are encoded in the same mRNA, therefore and might be expressed according to a so far unknown control regime (see section 3.4 bellow) To our knowledge no such a promoter driving transcription of an overlapped ORFs system had been identified before in the B3-like genome sequences or in other Mu-like phages described so far.

**Fig. 3.**
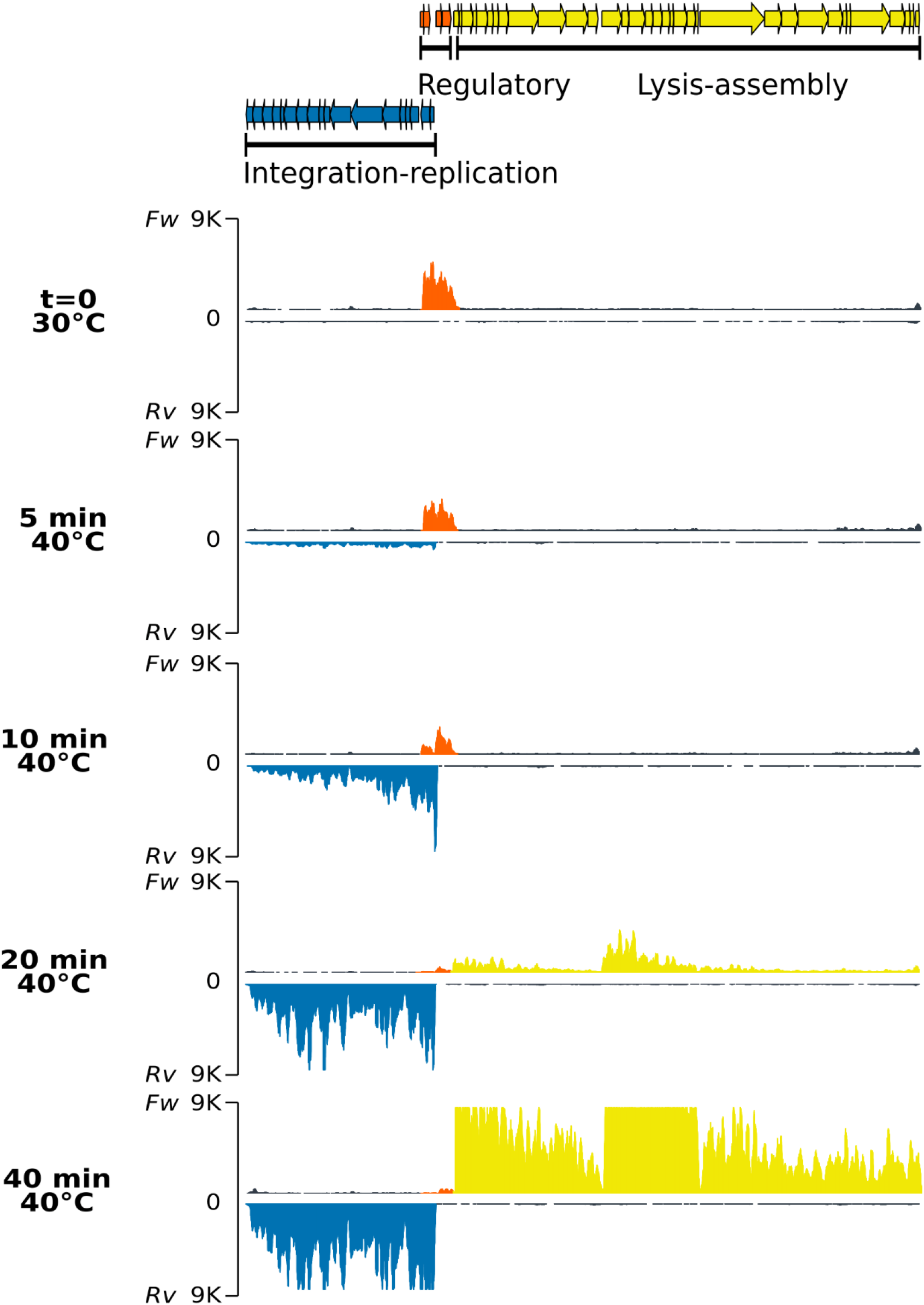
Strand-specific RNA-Seq of the phage genome after thermo-induction of lysogen PAO1(Fc02 *repc* ts). Samples of the lysogen culture grown at 30°C corresponds to the uninduced culture(t=0) and samples after up shifted to 40°C were drawn at the indicated times (see Fig.2). The RNA reads for each time sample are split in two, the upper trace corresponds to reads mapped to the forward (*Fw*) and the lower trace to reverse (*Rv*) strands for each pair. Note that the scales of reads are inverted relative to each other. The colors of the read-blocks correspond to the group of genes in the gene map at the top of the figure: orange, genome region transcribed during lysogeny at 30C°; blue, left-arm lytic cycle genes, and yellow, right-arm lytic cycle genes. The histogram for the reads was generated by *ggplot* with 9K reads maximum set (vertical axis) and map length in nucleotides (horizontal axis).

Transcription of the *ner*-like gene, mapped to the reverse strand, seems to rapidly ascend by 10 min. Gene *ner* in other phages encodes a protein that turns off *repc* transcription preventing RepC synthesis (Oppenheim et al., 2005, Hulo et al., 2015). Transcription of *ner*-like gene, by itself may antagonize transcription of sequences upstream the repressor gene. In spite this transcription reduction, the *repc-e4* mRNA remains in the cell for some time. We do not know whether this transcript was initiated from pC2 de novo, or it is a remnant of the RNA initiated at pC1.

At 10 min after up-shift of a lysogen culture to 40°C, pE promoter was activated with a burst of transcripts starting at the antirepresor gene *ner* and dwindling distally on the early lytic genes on the reverse strand of the phage DNA. The levels of the pE operon transcripts increased up to a maximum at 40 min, the last time recorded (Figure 3 and Supplementary Figure 4). Most of the ORFs in the pE operon are not assigned to any function except *transposase A*, *transposase B*, *gemA* and *mor* homologous genes. In phage Mu, *transposase A* participates in replicative transposition during the phage lytic cycle (Montaño et al., 2012) and *mor*, the last gene in the left operon, is an activator of genes in the middle operon (Mathee and Howe, 1990). Other genes in the left operon correspond to accessory genes in a plasticity genomic region ie., that could be present or not in other beetreviruses (Carballo-Ontiveros et al., 2020).

Right downstream to the pair of genes *repc-e*4 on the forward DNA strand three clusters of genes sequentially transcribed after thermal induction. The first cluster starts with the holin-endolysin-like genes followed by several DNA packaging genes (Figure 3 and Supplementary Figure 4). The first group of genes is barely transcribed at 20 min when the genes of the pE operon have nearly reached their maximum level of transcription, whereas the second cluster of genes shows incipient transcription. This cluster includes mainly virion structural genes and appears to be limited by a rho-independent transcription terminator (Supplementary Figure 5). The third and first cluster of genes on the forward strand show about synchronous levels of transcripts ending at 40 min (Figure 3 and Supplementary Figure 4). Some accessory genes are intercalated in each of the clusters. The gene expression pattern through induction was confirmed by differential expression analysis (Supplementary Figure 6).

### 3.3 Gene structure of the lysis-lysogeny regulatory region of bacteriophage Fc02

The regulatory region in the Fc02 genome follows the genetic configuration of the B3-like phages of *P. aeruginosa*. In fact, transposable phages of other bacteria, share with Fc02 the genetic structure of the regulatory region (Figure 4) (Stoddard and Howe, 1989, Fogg et al., 2011, Cazares et al., 2014). Repressor and anti-repressor genes are oriented in opposite directions separated by a non-coding DNA stretch containing the overlapping promoters historically named pRM or pR, respectively and pRE necessary for the establishment of repression (Spiegelman et al., 1972, Echols and Green, 1971, Hayes and Slavcev, 2005). In Fc02, the DNA stretch is rather short (106 bp), such that the sequences of the promoters pC2 and pE overlap by some nucleotides (Figure 4B). At 30°C transcription of the repressor gene appears to occur from the distal pC1 promoter (Figure 4A). However, from the forward transcription profile it appears that transcription from pC2 is active at 10min after induction whereas pC1 activity seems to fade away gradually after induction. The presence of pC1 is rather conserved among the *P. aeruginosa* phages of the B3-family indicating its essential role in the phage lysogenic cycle (Supplementary Figure 7).

**Figure 4.**
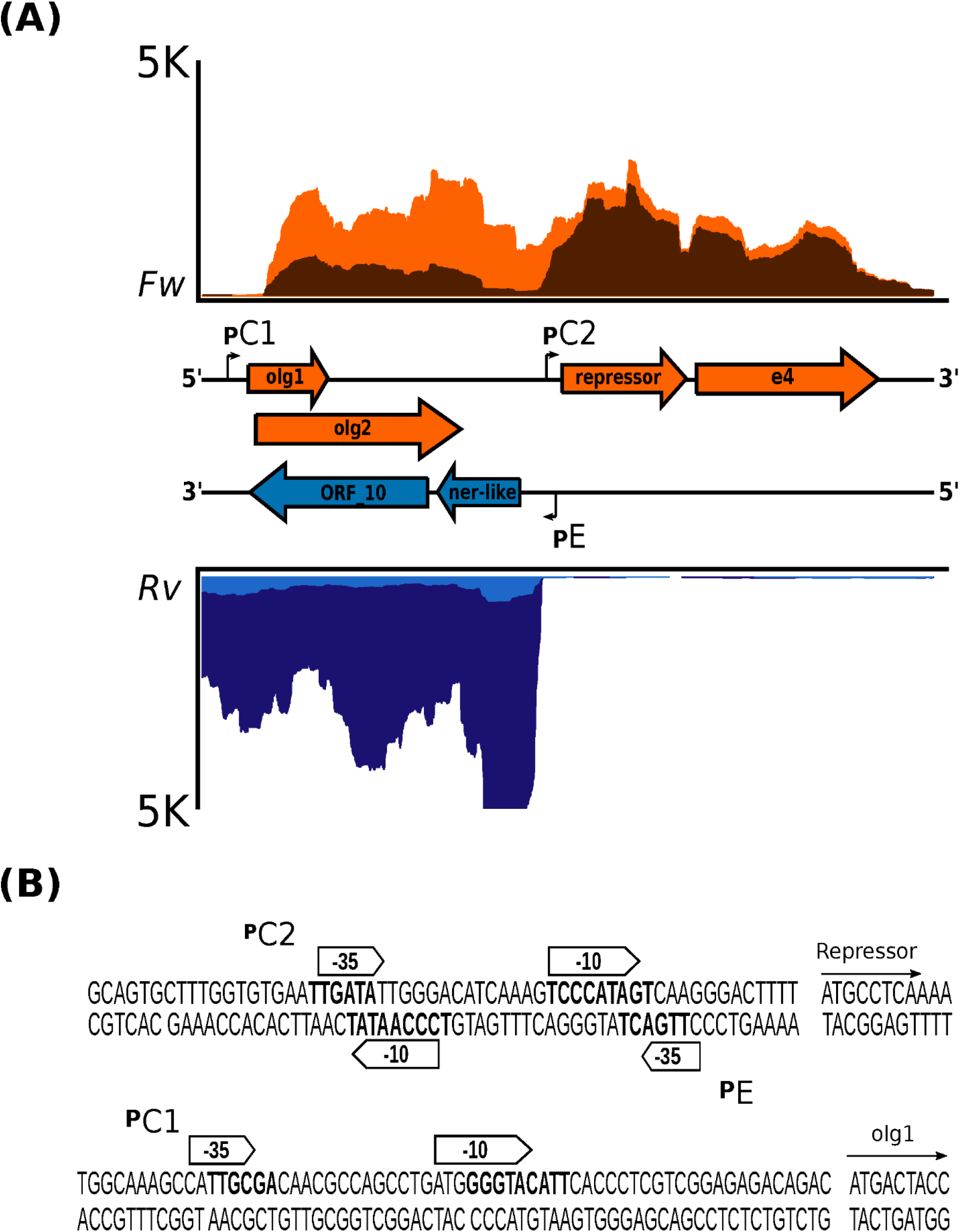
Genetic configuration and promoter sequences identified in the lysis-lysogeny regulation region of bacteriophage Fc02 genome. A. Blow up of the strand-specific RNA-Seq reads mapping to the regulation region in Fig. 3. Graphs in orange represent the reads coverage mapped to the forward strand (*Fw*) at 5 min (light orange) and 10 min after induction at 40°C (dark orange). Graphs in blue, inverted relative to the orange plots, denote the reads coverage of the reverse strand (*Rv*) at 5 min (light blue) and at 20 min (dark blue) after induction. In the middle, separating both graphs is presented a map to scale of the genes (colored arrows) and the location of the promoter sequences (angled arrows). The genes *olg1* and *olg2* are overlapped in the zero and +1 frames. The *e4* arrow is a putative accessory gene of unknown function. Two more short ORFs, located between *olg2* and pC2 are not shown. *In* both coverage graphs upper limit was +/- 5K reads (vertical axes). The horizontal axis represents the 1980 nucleotides length of the regulation region. B. Sequence detail of the putative promoters identified using BProm, SAPPHIRE and Promoter Prediction.

### 3.4 Identification of overlapping genes *olg1* and *olg2*

We then asked whether the transcript initiated at pC2 was translated despite being overlapped to gene 10 and partially to the *ner*-like gene in the reverse strand of the regulatory region (see Figure 4A). A search using software ORF Finder revealed the presence of a set of several ORFs, however we focused our attention on the two overlapped ORFs proximal to the promoter, *olg1* and *olg2*, inscribed within different frames in the forward strand of the regulatory region DNA. The left most ORF, *olg1*, is the closest to promoter pE and has a potential initiation AUG codon properly spaced from a putative SD sequence such that it is expected to be translated. The putative AUG initiation codon for the overlapped *olg2* was located 25 nucleotides downstream in the +1 frame. The final G of this AUG could be the first nucleotide of a stem-loop of a possible rho-independent terminator (Figure 5A). In this work the interactions of the elements in this complex configuration for the expression of *olg1* and *olg2* were not examined, however they are tantalizingly interesting to be tackled soon in the future.

**Figure 5.**
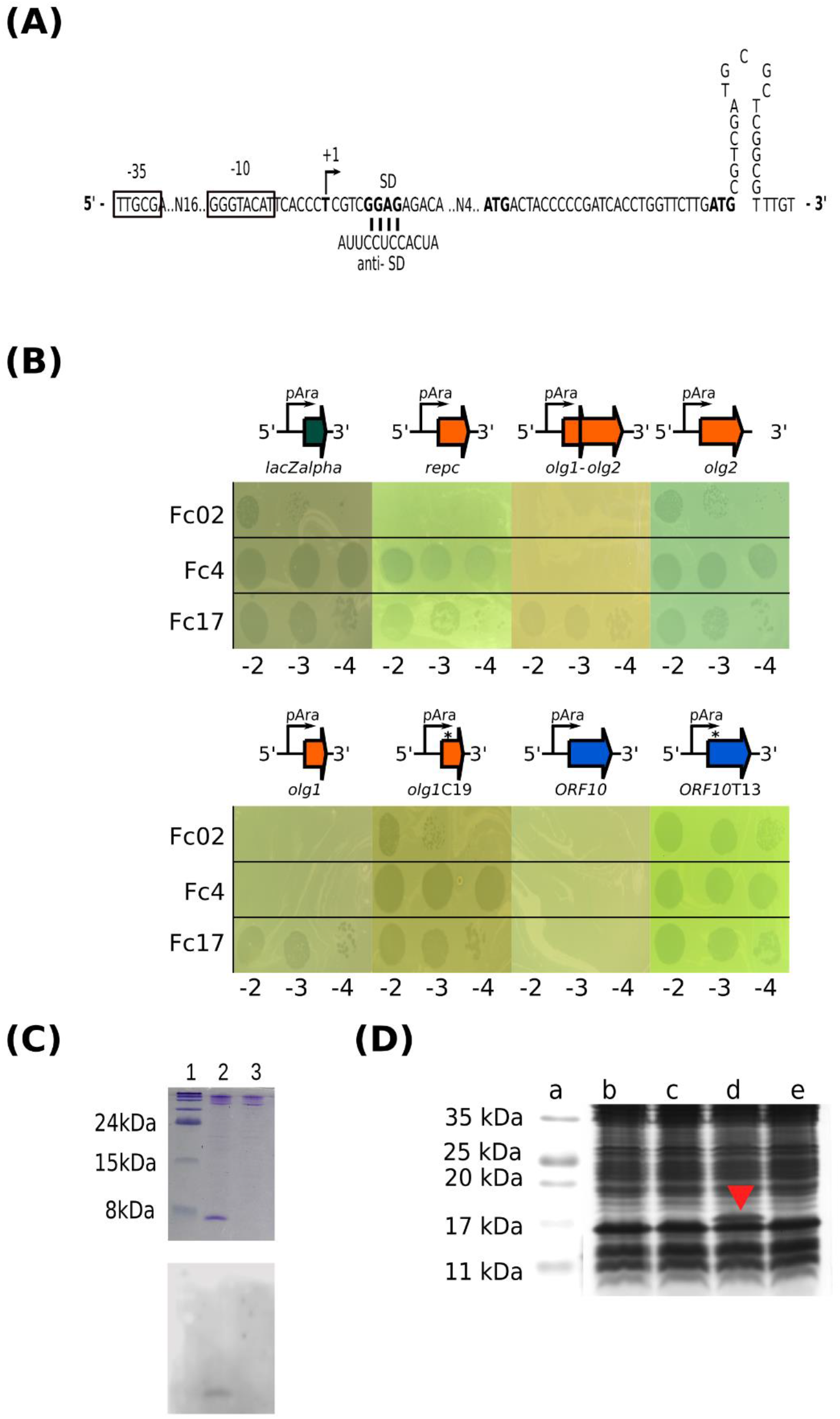
Exclusion activity of *olg1* and ORF10. A. Sequence detail of promoter and nucleotide sequence distance between the putative start codons from the overlapping genes *olg1* and *olg2*, indicating the stem-loop structure downstream *olg*2 start. B. Olg1 and Gp10 exclude other infectant phages. Plaque assays were carried out dropping dilutions of the indicated phages on 0.3% L-arabinose induced lawns of PAO1 harboring vector pHERD30T, or constructs pH*olg1*+*olg2*, pH*olg1*, pH*ORF10*, pH*olg1*C19 and *pHolg2*. C. Expression of *olg1*in PAO1. Samples were analyzed by Tricine-SDS-PAGE at 16% followed by Coomassie blue staining (upper panel) or Western immunoblot on a nitrocellulose membrane probed with anti-hexa His antibody (lower panel). Lane 1 BlueRay Prestained Protein Marker (Jena Bioscience); lane 2 Purified Olg1 protein after 3 h arabinose (0.3%) induction and Ni^2+^-affinity chromatography enrichment; lane 3 Purified Olg1 protein without induction after Ni^2+^-affinity chromatography. D. Expression of *ORF10* in PAO1. Samples were analyzed on Tricine-SDS-PAGE on 16% polyacrylamide gels followed by Coomassie blue staining. Lane a, Prestained Pritein Ladder (MaestroGen AccuRuler RGBPlus); lane b, uninduced ORF10; lane c, pHERD30T without inducer; lane d, ORF10 after 3h of arabinose induction; lane e, pHERD30T after 3h mock induction on arabinose. Red arrow-head indicates ORF10 protein band position in the gel.

As we do not understand how the proposed genes *olg1* and *olg2* are translated from the pE transcript, and how are they regulated in the lysogen, we cloned the respective ORFs, in constructs expected to express each of them separately (see M&M and Figure 5). Based on the proposed exclusion function assigned to the homologous gene 10 of phage Ps56, translatability and superinfection exclusion of *olg1* and *olg2* was assayed on transformed cells (Carballo et al., 2020). We assessed the exclusion of Fc4, Fc17 and Fc02 phages by cells expressing Olg1, Olg2 and ORF10 polypeptides from plasmid constructs. Under the used conditions Olg1 excluded Fc02 and Fc4, and ORF10 excluded all super-infecting phages assayed, but mutant versions of *olg*1 and ORF10 that generate premature stop codons, lost the exclusion function in transformed cells. In addition, we proved that *olg1* was in fact a translatable gene as the corresponding His-tagged protein was identified using Ni-resin enriched preparations and Western blots (see M&M and Figure 5C) and verified the ORF10 protein expression in a Commassie stained SDS-PAGE (Figure 5D). We were unable to observe expression of Olg2 or exclusion of the superinfecting phages, therefore *olg*2 function remains unknown.

## 4 Discussion

In this work we designed a phage-bacteria system to investigate the prophage transcription regime in the lysogen state and its transition to the different development lysis stages using strand-specific RNA-Seq technique. Concurrently with the induction of the Fc02 *repc* ts prophage, we observed changes in transcription of the bacterial genes. As expected, the upregulated genes shared between PAO1 and PAO1(Fc02 *repc* ts) after the first 5 minutes at 40°C, are involved in the heat-shock response(Chan et al., 2016). Many other changes associated with phage development were observed (Supplementary Figure 8 and Table S1 of Supplementary material).

Regarding phage transcription, we found that in the lysogen the repressor gene apparently is transcribed forward from a distal promoter pC1, but the transcript (ca. 1700 nucleotides) includes upstream genes. The upstream transcript region includes two overlapped genes, *olg*1 in frame +0 and *olg*2 in frame +1 of the DNA forward strand(Lèbre and Gascuel, 2017, Muñoz-Baena and Poon, 2022) and two smaller ORFs not studied in this work. Interestingly *olg1-olg2* overlapped region, in turn completely overlaps gene 10, and part of the 3’-end of *ner*-like gene, on the reverse strand. Remarkably, both *olg1* and gene 10 seem implicated in superinfection exclusion by Fc02 lysogen. Genes *ner* and10, in the early lysis left operon on the reverse strand, are transcribed from pE once the lytic response takes over after prophage induction and pC1is shut off (Figure 4). Although the regulatory switch issue has been studied in several temperate phage models, the expression and function of the ORFs in pC1 operon require to be investigated in Fc02 to be understood in detail.

In dsDNA viruses is frequent the incidence of overlapping genes, but these systems are often not well characterized. Either there is no proof that the genes involved are expressed, or their function is not clearly known(Muñoz-Baena and Poon, 2022, Schlub and Holmes, 2020). Three-gene overlapping configuration, like the one described here, is a very rare instance but is known to occur in phages (Fiddes and Godson, 1979).

On the forward strand of the phage regulatory region only a possible initiation AUG in *olg*1 seems to be located at the appropriate distance from a putative SD sequence to be translated (see Figure 5). We do not know yet the prophage regulation for *olg*1 and/or *olg2* mRNA translation. For translation of *olg2* it is possible that an attenuation mechanism might be involved because in a sequence model of the transcript, an apparent stem-loop in the mRNA sequester the possible initiation AUG of *olg*2 located 25 n downstream *olg*1 initiation codon. Other models of translational control could be frameshifting. The presence of a CCCCCG, a possible slippery sequence followed by a stable secondary RNA structure downstream, could function as frameshift signals(Scherbakov and Garber, 2000). Expression of *olg*1 from an ad hoc construct driven by a plasmid promoter shows exclusion properties because transformants expressing Olg1 polypeptide prevent superinfection by a set of test phages (see Figure 5B). Therefore, we propose to rename *olg1* as *sie1* (for superinfection exclusion) following an acronym coined almost 30 years ago (Hofer et al., 1995, Ranade and Poteete, 1993). A BLASTp search of *olg*1 ORF showed that most of its predicted amino acid sequence matches a putative conserved domain present in a DEDDh exonuclease family, that could be associated with the exclusion activity at DNA level.

The reverse DNA strand, opposite to the *olg1-olg2* forward strand, contains gene 10 in beetre phages (Fig. 4). In phage Ps56, akin to Fc02, it has been reported that the gene 10 homologue grants exclusion properties to the lysogenic cells (Carballo-Ontiveros et al., 2020). Remarkably, overexpression of ORF10 in PAO1 also generates exclusion activity against temperate phages.

However, the exclusion mechanisms for the two genes might be different; gene10 of phage Ps56 seems to exclude by preventing injection of the superinfecting phage DNA (Carballo-Ontiveros et al. 2020). This may explain why ORF10 can exclude Fc17 but Olg1 cannot. We do not know much about Fc17 other than it is a temperate phage with a repressor specificity different from that of Fc02 repressor (Figure 5B).

Two overlapped genes expressing proteins with similar functions, in one direction throughout lysogeny and in the opposite direction during lytic development, summons to speculate about the possibility of a new kind of forward and reverse iso-functional DNA module. Whether they exclude at different steps of superinfection remains to be explored.

## 6 Author Contributions

Design of the work IRS, GG. Data collection and experiments IRS, MMC. Data analysis and interpretation IRS, GG. Drafting the article IRS, GG. Critical revision of the article IRS, GG. Final approval of the version to be published IRS, MMC, GG.

## 8 Acknowledgments

We thank Maria Guadalupe Aguilar Gonzalez (Unidad de Acidos Nucleicos, DGBM) and Dulce Delgadillo Alvarez (LaNSE-CINVESTAV) for technical support in sequencing the cloned ORFs constructs, Dr. David Romero Camarena, Dra. Sofía Carolina Martínez Absalón and M. Luis Jaramillo, for the bioinformatic support. The pHERD plasmids were provided by Ryan Withers.

## 10 Data Availability Statement

The datasets generated for this study can be found in the Short Reads Archive (SRA) database with accession number: PRJNA872573 https://www.ncbi.nlm.nih.gov/sra/PRJNA872573

