## Supplementary Figures for "Transcriptional analysis in bacteriophage Fc02 of *Pseudomonas aeruginosa* revealed two overlapping genes with exclusion activity"

### *Supplementary Material*

#### 1 Supplementary Figures

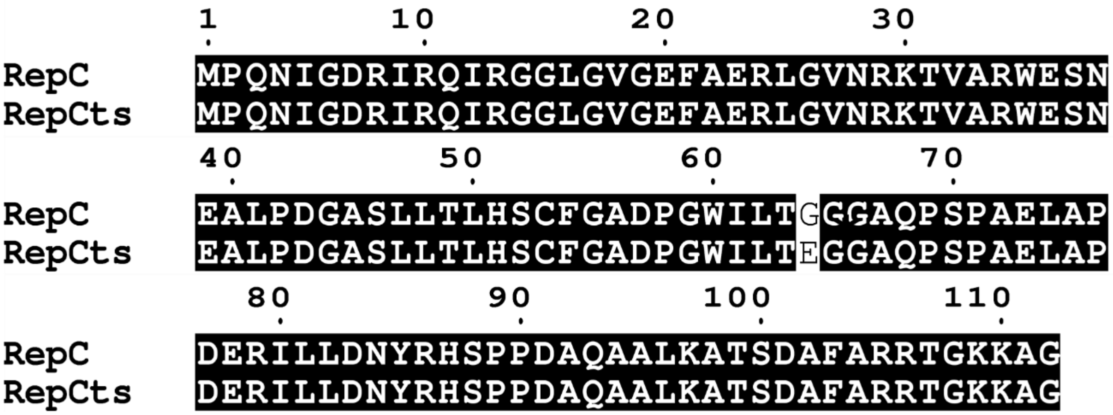


**Supplementary Figure 1. Fc02 *repc* ts repressor mutation.** Amino acid sequence alignment of the wild type of repressor protein and the mutated RepC ts repressor. Alignment was done according (Robert and Gouet, 2014, Sievers et al., 2011).

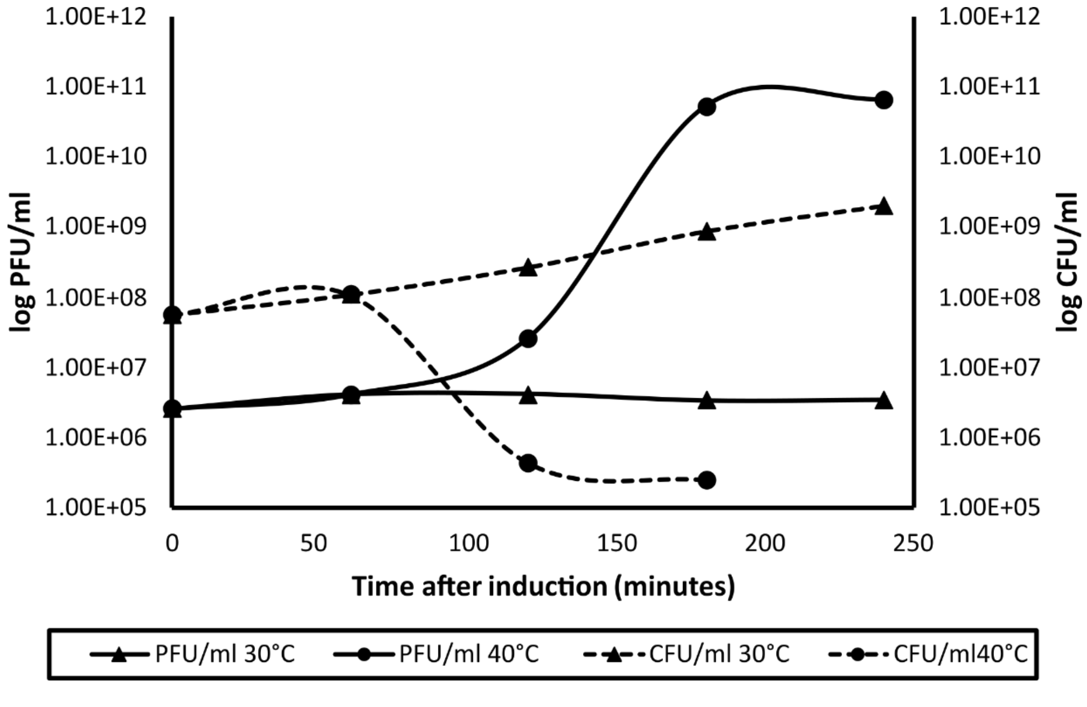


**Supplementary Figure 2. Time course of PAO1(Fc02 *repc* ts) growth at 30°C and after up shift to 40°C.** Measures were done on LB agar medium counting colony forming and plaque forming units (CFU and PFU) on a lawn of strain PAO1.

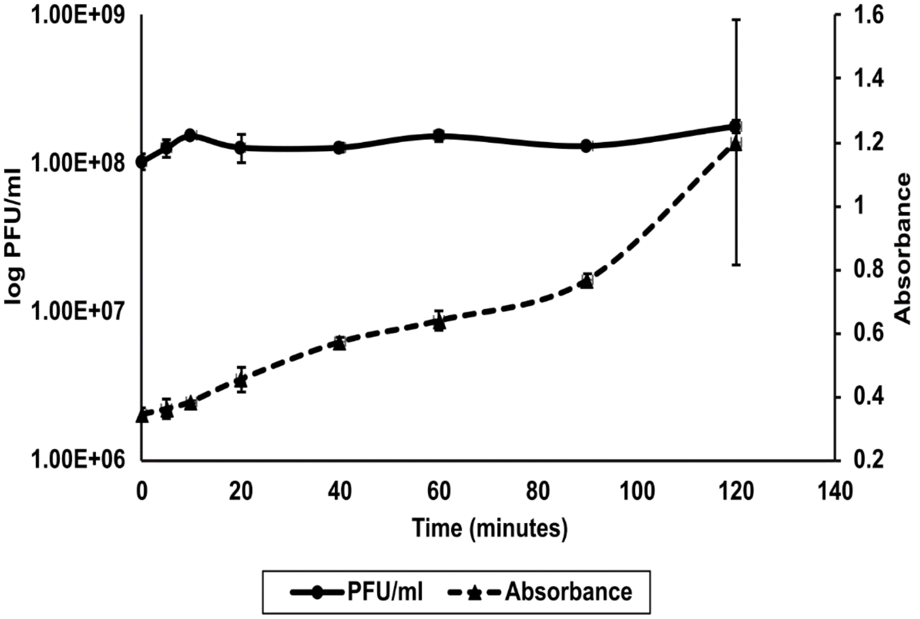


**Supplementary Figure 3. Phage release at 30°C and 40°C.** Time course of the lysogen PAO1(Fc02) growth at 30°C and shifted to 40°Cat time 0. Plaque forming units measured on a lawn of strain PAO1 and absorbance was determined at $OD_{600nm}$.

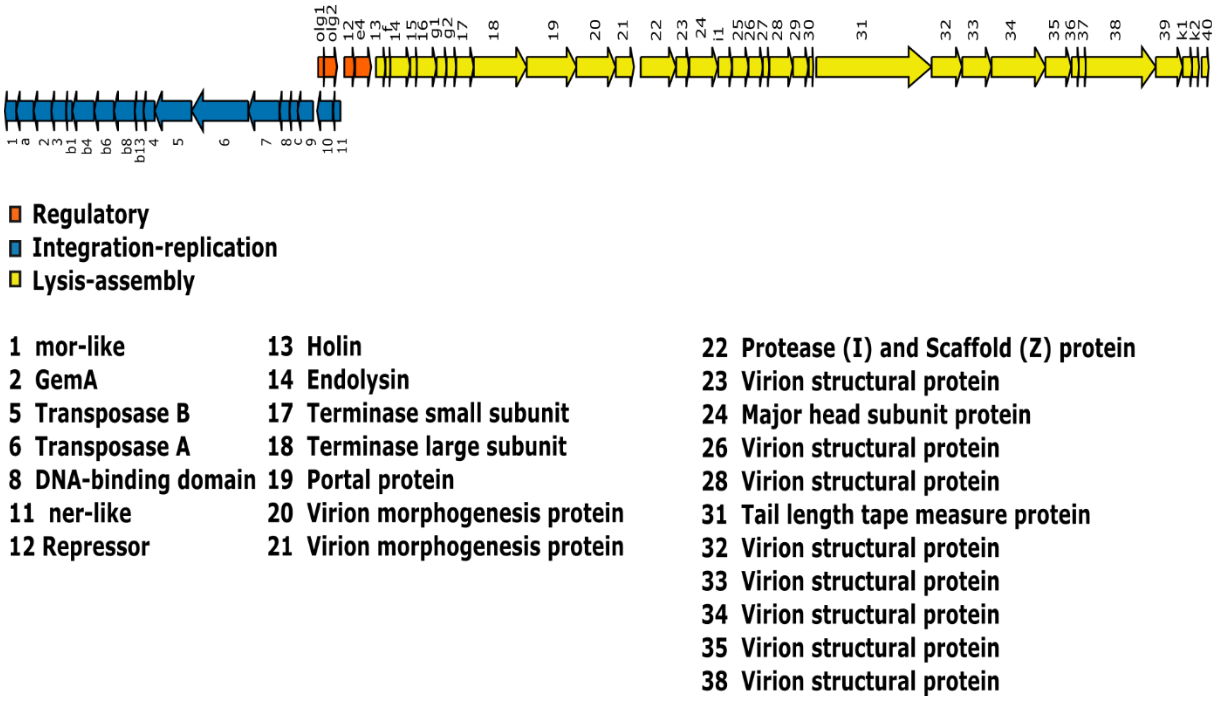


**Supplementary Figure 4. Fc02 genome map.** Genes are indicated as numbers or letters (for accessory genes).The genes with assigned function are enlisted at the bottom of the figure.(Carballo-Ontiveros et al., 2020)

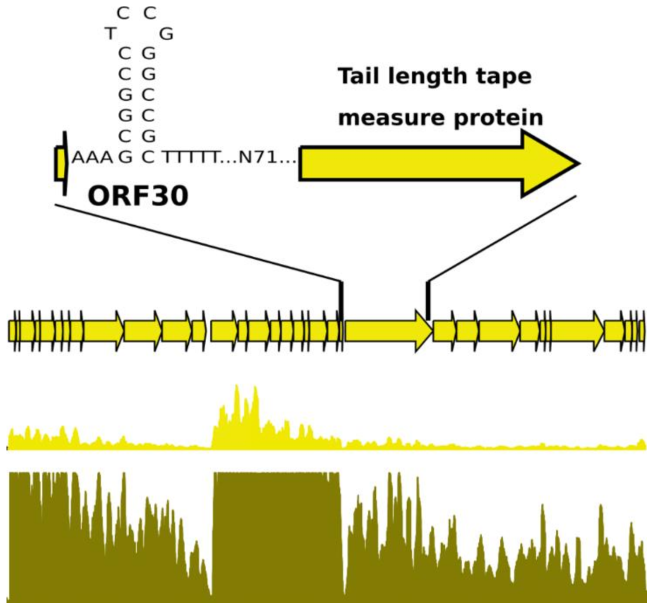


**Supplementary Figure 5. Close up of the late genes' transcription.** Graphs represent the reads coverage mapped to the forward strand at 20 min (yellow) and 40 min (dark yellow) after induction at 42°C. The maximum reads in the vertical axis is limited to 9K. The putative terminator between right lysis genes 30 and 31 is indicated above the upper map.

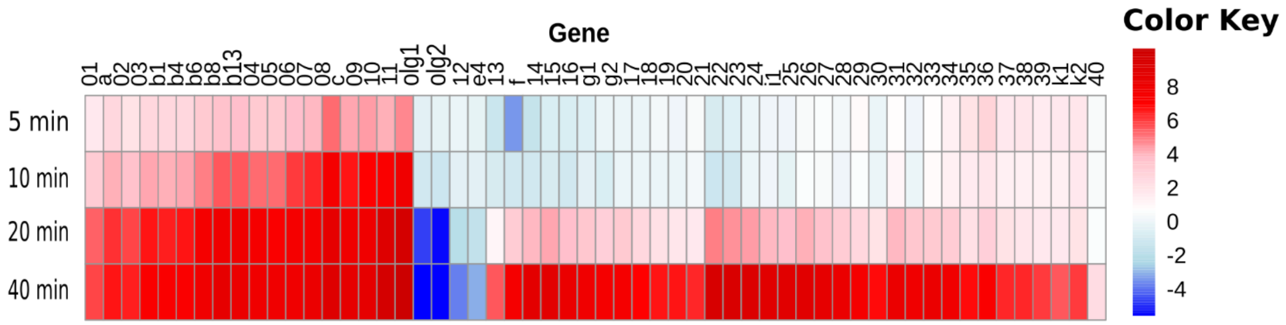


**Supplementary Figure 6. Heatmap of the Fc02 genes after induction. T**ranscription intensities of phage Fc02*repc* ts genes induced in a PAO1 lysogen relative to an uninduced sample (t=0) are represented as two-fold change using software DeSeq2 (Love et al., 2014). Color key is indicated on the right hand of the figure. Blue color density indicates transcription below that at time 0 at the indicated times and red color density transcription above time 0 after induction. The corresponding Fc02 genes are indicated above the columns.

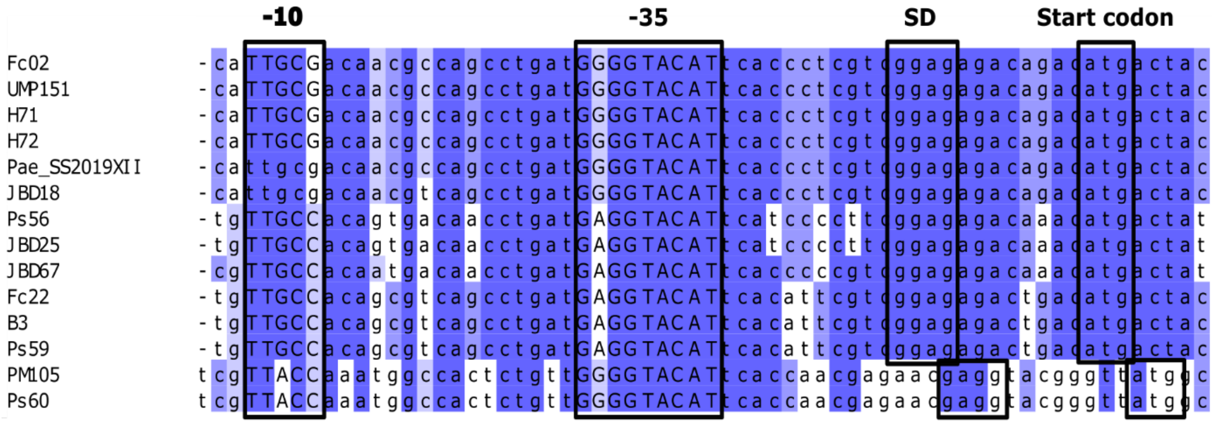


**Supplementary Figure 7. Alignment of the promoter region pC1 in genomes of beetreviruses.** Phages considered are those filed as free virions that infect *P.aeruginosa*. Not prophages were included (Gouy et al., 2010, Waterhouse et al., 2009).

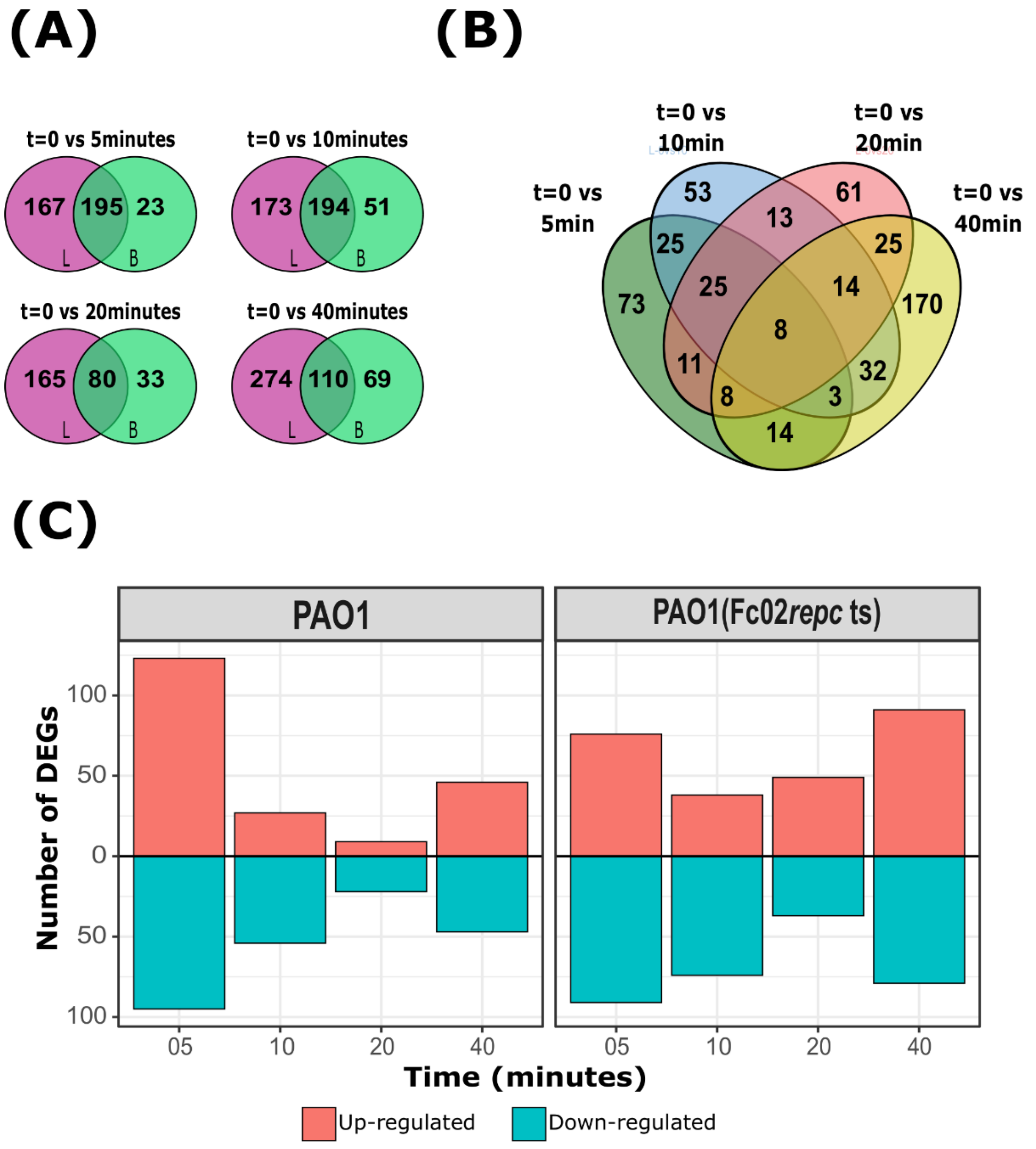


**Supplementary Figure 8. Differentially transcribed PAO1 genes. A.** Venn diagrams comparing PAO1and PAO1(Fc02 *repc* ts). Circles represent the total number of genes differentially transcribed in each time after up shifting temperature at 40°C relative to time 0 in the PAO1 strain, green circles(B), and in PAO1(Fc02 *repc* ts) violet circles(L). B Venn diagram of the total of differentially transcribed genes in PAO1(Fc02 *repc* ts) at 5-, 10-, 20- and 40-minutes relative to time 0. C. Bar plots indicating the total of upregulated and downregulated genes in both strains PAO1 and PAO1(Fc02 *repc* ts).
